# PfHT1 controls glucose uptake in malaria parasite: molecular dynamics study in plasma membrane like environment reveals ligand induced conformational changes trigger closed conformation and alters binding pocket geometry

**DOI:** 10.1101/2024.05.01.592028

**Authors:** Manisha Ganguly, Hriddhi Maitra, Amlan Roychowdhury, Ditipriya Hazra

## Abstract

*Plasmodium falciparum* hexose transporter 1 protein or PfHT1 is the major glucose transporter of the parasite and indispensable for its survival at the blood stage of infection. PfHT1 transports the hexose sugar to meet the energy demand of the parasite. Studying the mechanism of transport and designing structure specific inhibitors against PfHT1 is an intelligent strategy to kill malaria parasite by starving it out because at the blood stage the sole carbon source of *Plasmodium falciparum* is glucose. In this study the conformational dynamics of PfHT1 has been studied in detail in the apo and holo (inhibitor bound) form using Molecular Dynamics Simulation. Which reveals that PfHT1 undergoes ligand induced closed state if compared with the apo form. The geometry of ligand binding pocket also shifts from the apo form in the presence of inhibitors. A de novo drug designing approach based on the skeleton of leads obtained from screening nearly 4500 compounds, has produced inhibitors of PfHT1 with higher specificity and affinity. The drug screening data as well as the conformational dynamics study was validated using Molecular Dynamics Simulation platform where a near physiological atmosphere was created for PfHT1 by constructing a lipid phase (phospholipid bilayer) sandwiched between aqueous phases (mimics extracellular and cytosolic polar environment).

## INTRODUCTION

Malaria, has prevailed as an endemic in African and South-east Asian countries, despite aggressive efforts to eradicate the disease, in 2022 approximately 249 million people were reported to have contracted the disease (Milner, 2018). Malaria remains a persistent health challenge in underdeveloped countries, particularly in sub-Saharan Africa and Southeast Asia. The emergence of drug-resistant Plasmodium strains, including resistance to chloroquine and artemisinin derivatives, exacerbates the situation (Takala-Harrison & Laufer, 2015). The perilous disease transmitted by mosquito-borne *Plasmodium falciparum* parasites, exhibits a complex mode of action within the human host. The parasites undergo a series of developmental phases, including rings, trophozoites, and schizonts, during the asexual intraerythrocytic cycle. At the schizont stage, the parasites rupture red blood cells (RBCs), releasing merozoites that promptly infect new host cells, perpetuating the infection (Mawson, 2013). At the blood stage the pathogen relies on glucose as its sole carbon source (Kirk, 1996, 2001). In *P. falciparum* the major glucose transporter protein PfHT1 (*P. falciparum* hexose transporter 1), is encoded by the gene PF3D7_02047001, without any close paralogue (Slavic et al., 2011; Woodrow et al., 1999) and this protein was shown to be essential for survival of the blood stage parasite (Blume et al., 2011). Glucose transporters play a pivotal role in cellular metabolism by facilitating the uptake of glucose, a fundamental energy source. Among these transporters, the human glucose transporter 1 (GLUT1) holds particular significance, as it serves as the primary glucose transporter in erythrocytes (Deng et al., 2014). Its crucial role in ensuring glucose supply to various cells underscores its importance in maintaining cellular energy homeostasis. PfHT1, analogous to human GLUT1, serves as the transporter system in *Plasmodium falciparum*, causative agent of malaria. Developing a PfHT1 specific inhibitor that can interrupt glucose transport in malaria parasite but hardly affecting the human glucose transporters would be an effective strategy to create a glucose starved condition which will have a lethal effect on the parasite.

PfHT1, shares only 27.4% sequence similarity with human GLUT1. It has also been reported that PfHT1 exhibits a distinctive feature—an additional pocket at the inhibitor binding site, connected by a narrow channel. The pocket is highly hydrophobic in PfHT1, in contrast to the hydrophilic nature observed in hGLUT1(Jiang et al., 2020b). This structural dissimilarity presents a valuable opportunity for the development of a PfHT1-specific inhibitor through structure-guided drug design suggesting the potential for selective inhibition and an attractive target for antimalarial drug development. Previous studies have identified a small molecule, compound C3361, capable of selectively inhibiting PfHT1 and suppressing the in vitro proliferation of *P. falciparum* (Jiang et al., 2020a).

Finding a novel drug against the malaria parasite needs more extensive research which includes studying protein ligand interaction in plasma membrane environment, brute force screening and structure guided rational drug designing approach. In this study, we aim to leverage the structural difference between of PfHT1 to GLUT1, coupled with the identified structural distinctions, to develop a targeted antimalarial drug. Our approach involves an unbiased brute-force screening of 4500 molecules, followed by molecular dynamic simulations in a near physiological environment which mimics plasma membrane (Figure 1) to identify potent lead compounds with strong binding interactions with PfHT1 and studying the conformational changes occurred in PfHT1 due to ligand binding. To enhance drug specificity, we employ structure-based drug design to manipulate the molecular skeletons of lead compounds, ensuring druggability, ease of synthesis, and minimal interaction with human GLUT1.

**Figure 1:**
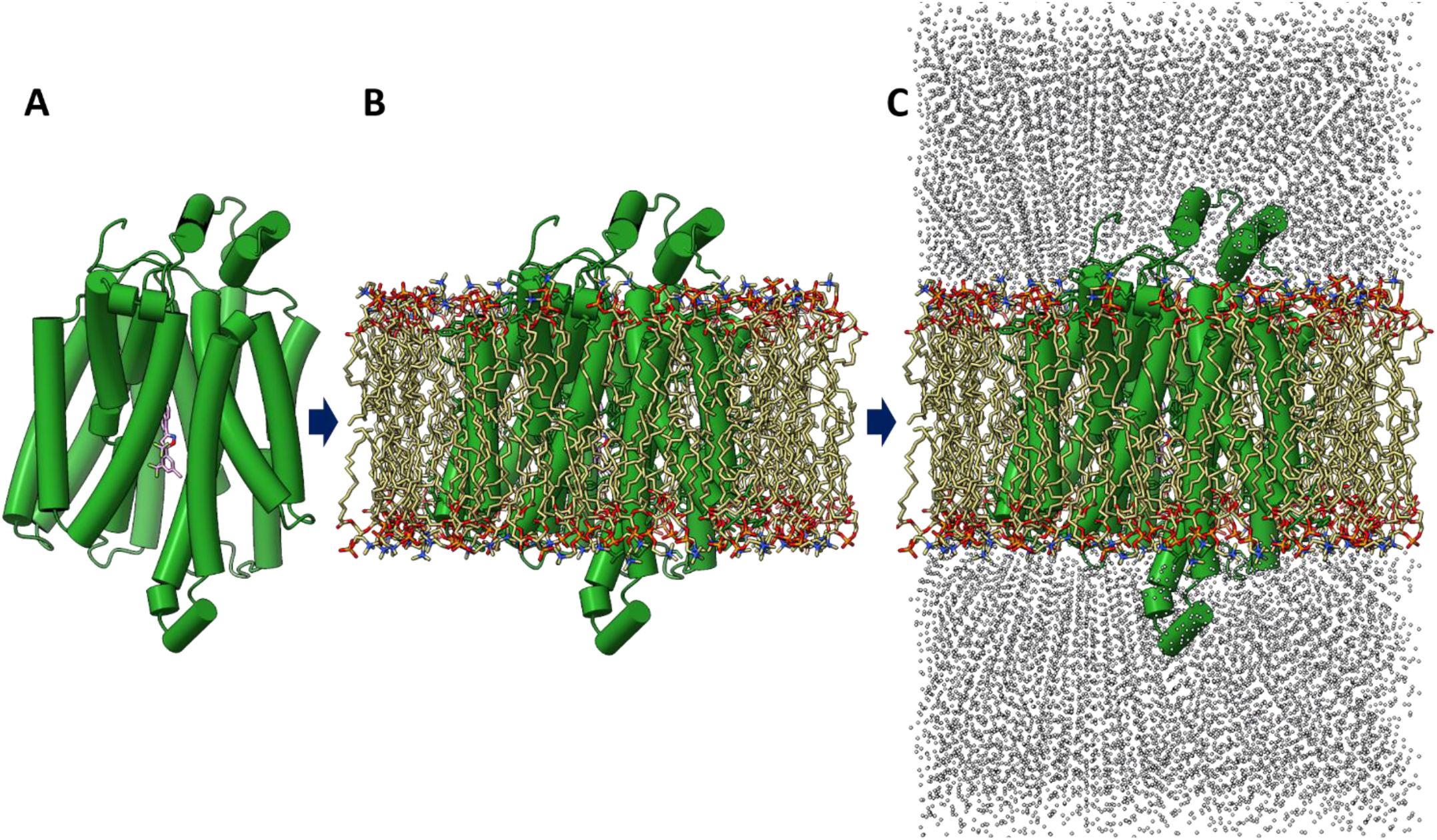
Graphical representation of the workflow adapted to create the physiological environment for MD simulation of PfHT1, (A) Protein alone, (B) Protein within the phospholipid bilayer, (C) Final assembly for MD simulation where the protein in phospholipid bilayer is sandwiched between the upper and lower aqueous environment.

## RESULTS AND DISCUSSION

### Molecular Docking Analysis for PFHT1 Inhibitor Discovery

In this investigation, a high throughput screening of around 4500 compounds, comprising commercially available compounds and FDA-approved drugs with accessible 3D conformers, was conducted using molecular docking with Autodock Vina (ADV) (Trott & Olson, 2010). The primary objective was to identify potential leads for rational drug design targeting PfHT1. To validate the efficacy of the docking protocol employed in this screening, the receptor was docked with C3361. The resulting docking score, expressed in kcal/mol, was recorded as ADV energy, yielding a value of -6.5 kcal/mol. The docking protocol was validated by superimposing docked C3361 with the crystal structure. The brute force screening results were initially ranked according to ADV energy, with the docking of C3361 (−6.5 kcal/mol) serving as a reference cutoff. Brute force screening gave over three thousand molecules above the cutoff. The top four compounds, Amentoflavone (−12.4 kcal/mol), Esafoxolaner (−11.8 kcal/mol), Iberdomide (−11.2 kcal/mol) and Veratramine (−10.8 kcal/mol), were subjected to molecular dynamics simulation. A tabulated representation of the docking scores for the top 4 compounds, along with their chemical structures and assigned three-letter codes for reference, is presented in Figure 2. These three letter codes are utilized for subsequent references in the manuscript.

**Figure 2:**
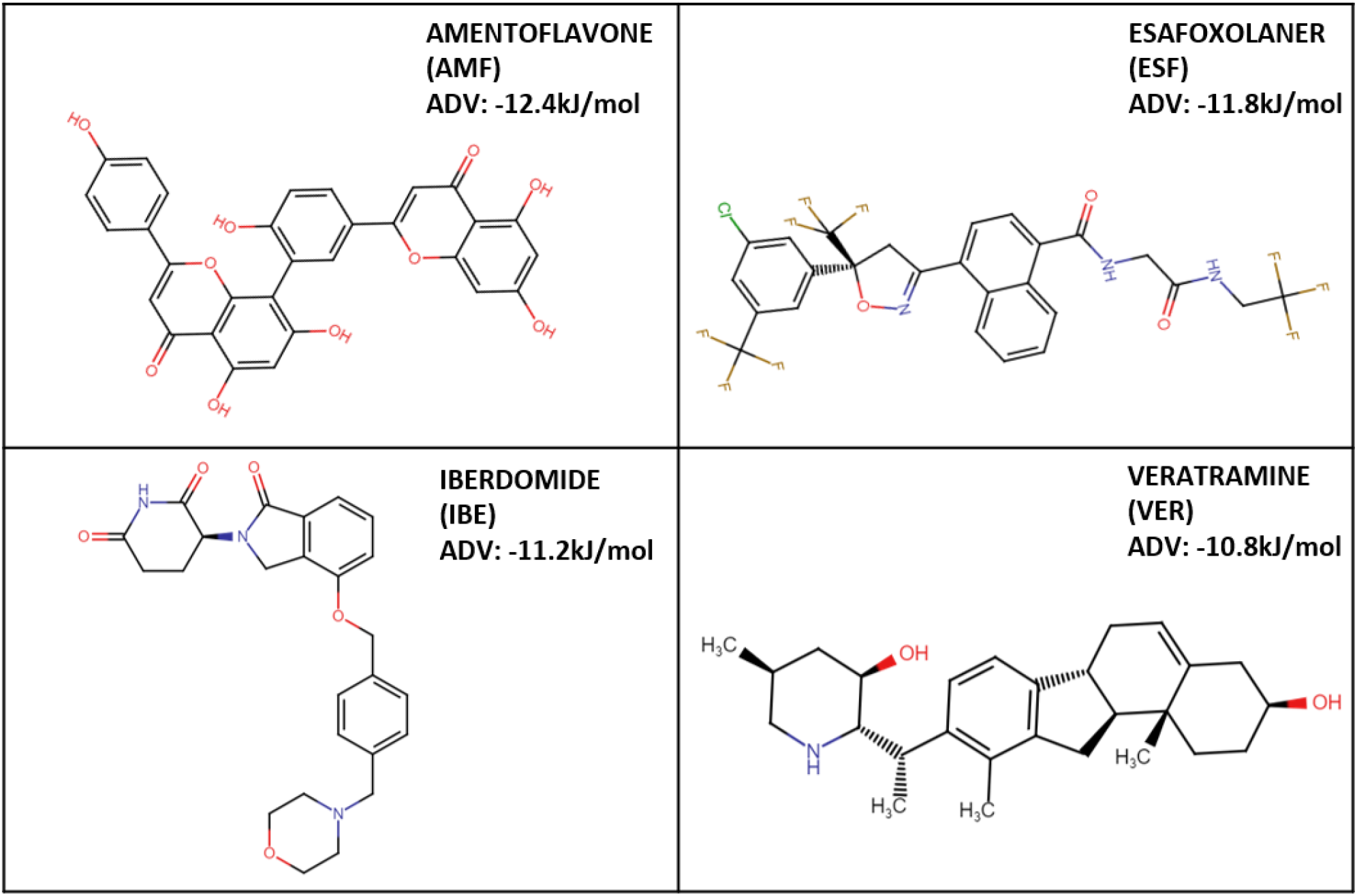
Name, chemical structure, and the three-letter code of the top 4 molecules according to ADV binding energy.

### ADME and Toxicity Assessment

The evaluation of druggability and toxicity for the top 4 compounds, identified based on ADV energy, were assessed using SWISS-ADME (Daina et al., 2017). Druggability assessment adhered to Lipinski’s rule of five, considering parameters such as molecular weight, n-Octanol partition coefficient, and the count of hydrogen bond donors and acceptors. To further scrutinize the safety profile of these compounds, toxicity analysis was performed using the Pro Tox-II server (Banerjee et al., 2018). The outcomes indicated that the molecules belonged to class 4. It is important to note that higher class values correspond to lower toxicity, where class 1 signifies extreme toxicity, and class 6 denotes non-toxicity. The ADME and toxicity results are tabulated in Table1.

**Table 1:** ADME and Toxicity assessment for AMF. ESF, IBE, VER.

| Parameters | AMF | ESF | IBE | VER |
| --- | --- | --- | --- | --- |
| MW | 538.46 | 625.87 | 449.5 | 409.6 |
| Acceptable range | $\leq 500$ | $\leq 500$ | $\leq 500$ | $\leq 500$ |
| NHBA | 10 | 11 | 6 | 3 |
| Acceptable range | $\leq 10$ | $\leq 10$ | $\leq 10$ | $\leq 10$ |
| NHBD | 6 | 2 | 1 | 3 |
| Acceptable range | $\leq 5$ | $\leq 5$ | $\leq 5$ | $\leq 5$ |
| MR | 146.97 | 134.5 | 131.49 | 127.88 |
| Acceptable range | 40–130 | 40–130 | 40–130 | 40–130 |
| ilogp | 3.06 | 3.65 | 3.08 | 4.12 |
| Acceptable range | $\leq 5$ | $\leq 5$ | $\leq 5$ | $\leq 5$ |
| ProTox class | 4 | 4 | 4 | 4 |

### Molecular Dynamics Simulation and Binding Free Energy Calculation

The top 4 compounds identified based on ADV energy (Figure 2) were subjected to Molecular Dynamics Simulation (MD simulation) to evaluate their dynamic behaviour and ascertain stronger interactions. Additionally, binding free energy calculations were performed to refine the selection of potential leads.

### Simulation Metrics

To assess the stability of the protein-ligand complexes, the following parameters were analysed, Root Mean Square Deviation (RMSD), Root Mean Square Fluctuation (RMSF), Solvent Accessible Surface Area (SASA), and Radius of Gyration (Rg). The average RMSD, RMSF, SASA and Rg are tabulated in Table 2.

**Table 2:**
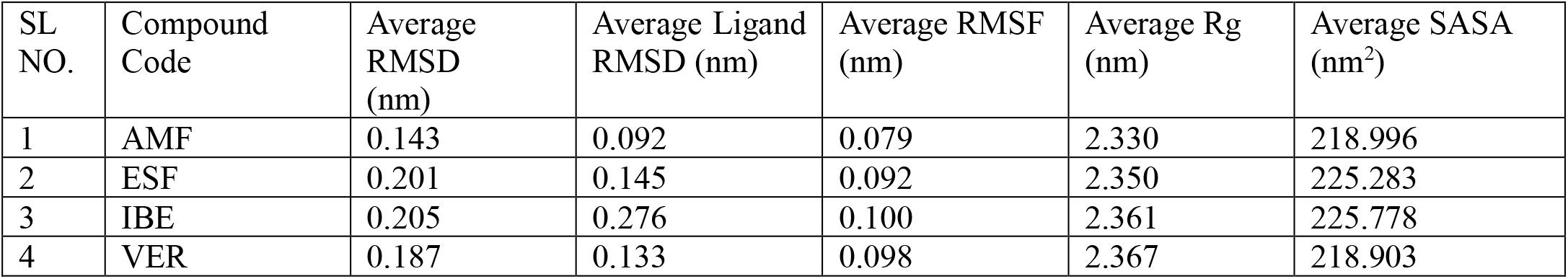
Simulation metrics for AMF, ESF, IBE, VER.

| SL NO. | Compound Code | Average RMSD (nm) | Average Ligand RMSD (nm) | Average RMSF (nm) | Average Rg (nm) | Average SASA (nm<sup>2</sup>) |
| --- | --- | --- | --- | --- | --- | --- |
| 1 | AMF | 0.143 | 0.092 | 0.079 | 2.330 | 218.996 |
| 2 | ESF | 0.201 | 0.145 | 0.092 | 2.350 | 225.283 |
| 3 | IBE | 0.205 | 0.276 | 0.100 | 2.361 | 225.778 |
| 4 | VER | 0.187 | 0.133 | 0.098 | 2.367 | 218.903 |

#### RMSD

The average RMSD of the apo-protein was 0.22 nm. Protein-inhibitor complexes exhibited comparable average RMSD (0.14 - 0.20 nm) (Figure 3A).

**Figure 3:**
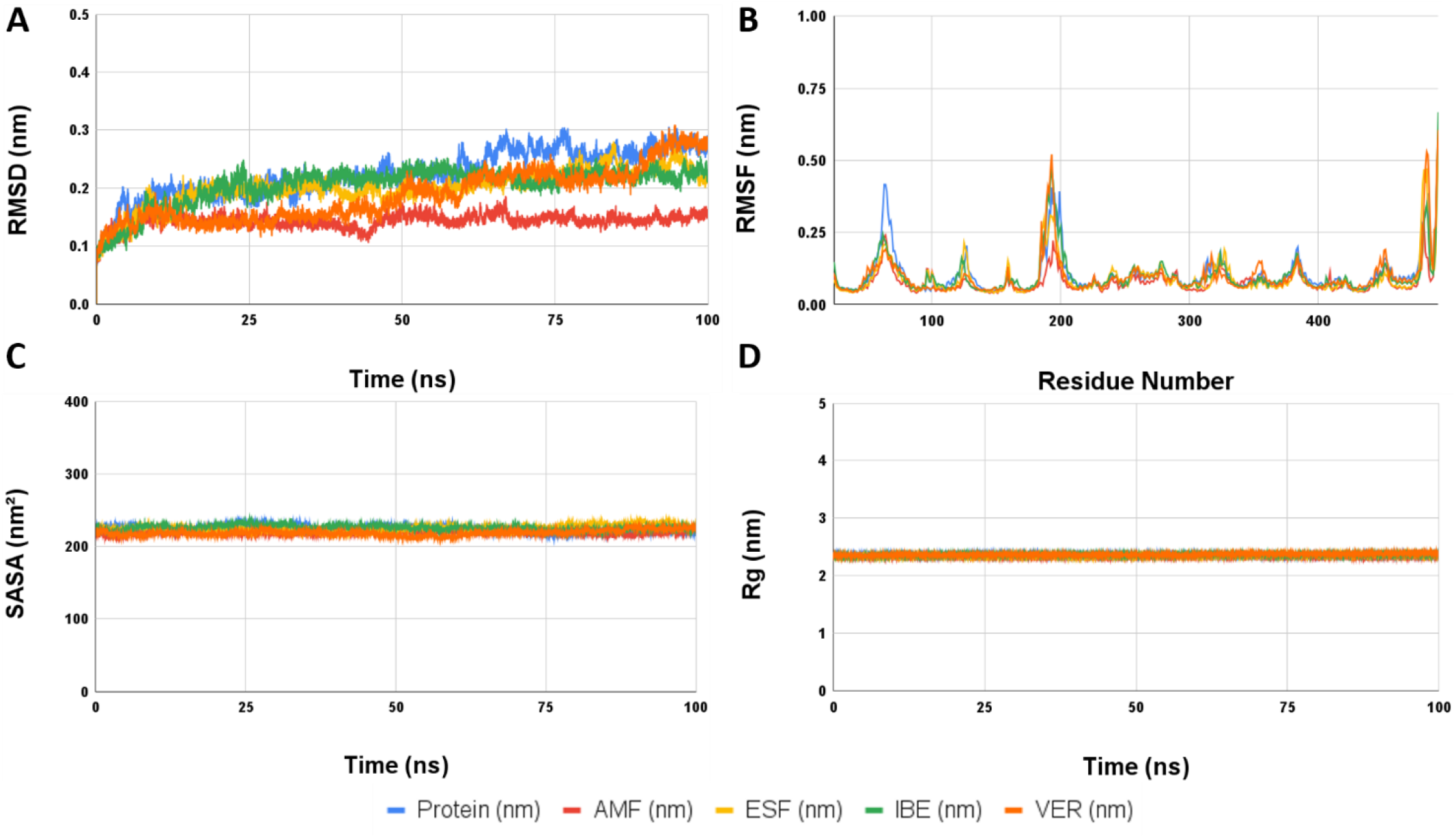
Molecular dynamics metrics of PfHT1 in apo and in the inhibitor bound form. The inhibitor bound protein complexes and the protein alone are shown in different colours. (A) The RMSD (nm) of backbone vs time in ns(B) backbone RMSF (nm) against the amino residue number, (C) solvent accessible surface area (SASA) in nm2 vs time (ns) and (D) Radius of gyration (Rg) in nm vs time in ns.

#### RMSF

Reflecting the flexibility of amino acid residues, RMSF values ranged from 0.079 nm to 0.100 nm for protein-inhibitor complexes, with an average of 0.104 nm for the protein alone (Figure 3B).

#### SASA

The solvent-accessible surface area (SASA) varied from 215.17 nm^2^ to 225.77 nm^2^ for inhibitor-bound complexes, with the protein alone having a SASA of 224.26 nm^2^ (Figure 3C).

#### Rg

The average Rg for the protein was 2.36 nm, with inhibitor-bound complexes maintaining a comparable range of 2.33 - 2.36 nm. This consistency indicated the overall structural compactness throughout the simulation (Figure 3D).

These simulation metrics collectively provide insights into the dynamic behaviour and structural integrity of the protein-ligand complexes, aiding in the identification of compounds with favourable interactions for further consideration.

### MM-PBSA Calculation

Utilizing the last 20 ns trajectories of MD simulation, the binding free energy (ΔG) was calculated for the protein-small molecule complexes of ESF, AMF, IBE, VER. All 4 demonstrated highly favourable binding (ΔG <-95 kJ/mol) calculated using the molecular mechanism Poisson Boltzman Surface Area (MM-PBSA) method (Figure 4A)(Kumari et al., 2014). Furthermore, AMF and ESF, showed significantly high binding energy of - 137kJ/mol and -174 kJ/mol hinting towards a higher affinity of these compounds for PfHT1.

**Figure 4:**
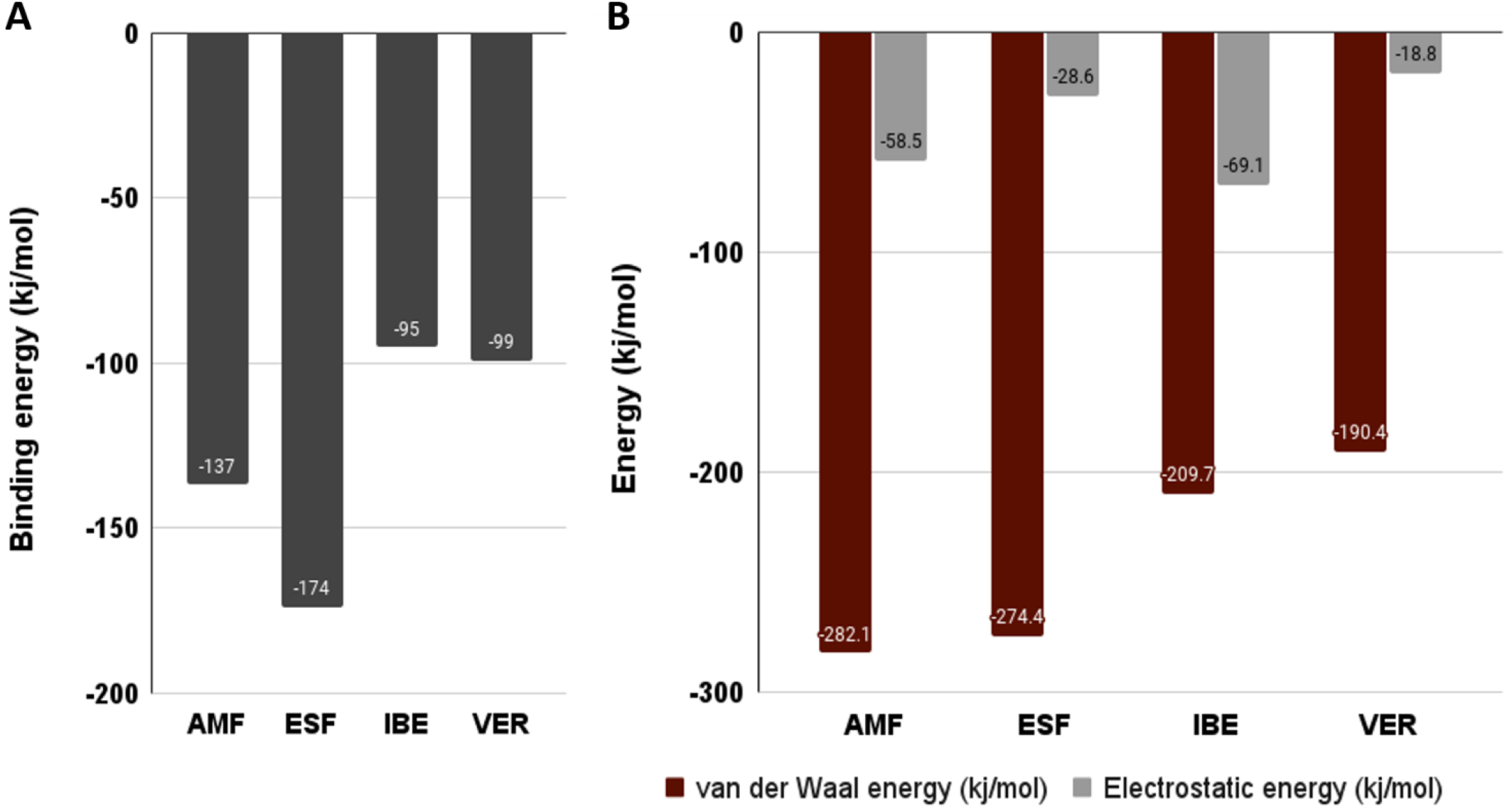
(A) Binding free energy (ΔG) for top 4, PfHT1-inhibitor complex expressed in kJ/mol. (B) The relative contribution of van der Waal’s energy and electrostatic energy in binding (expressed in kJ/mol).

### Protein-Ligand Interaction Analysis

To elucidate the nature of interactions governing the dynamics of protein-ligand complexes during simulation, Ligplot analysis was employed (Wallace et al., 1995). Coordinated data of each inhibitor-protein complex, extracted every 20 ns, were scrutinized to delineate the contribution of hydrogen bonds and hydrophobic interactions. Figure 5 illustrates the predominant role of hydrophobic interactions in steering the protein-ligand dynamics. Lennard-Jones potential and Coulombic short-range interaction energy analysis further affirmed the prevalence of van der Waals interactions over electrostatic forces (Figure 4B). The atomic level interaction of PfHT1 with AMP and ESF along with the binding pose are detailed in Figure 6.

**Figure 5:**
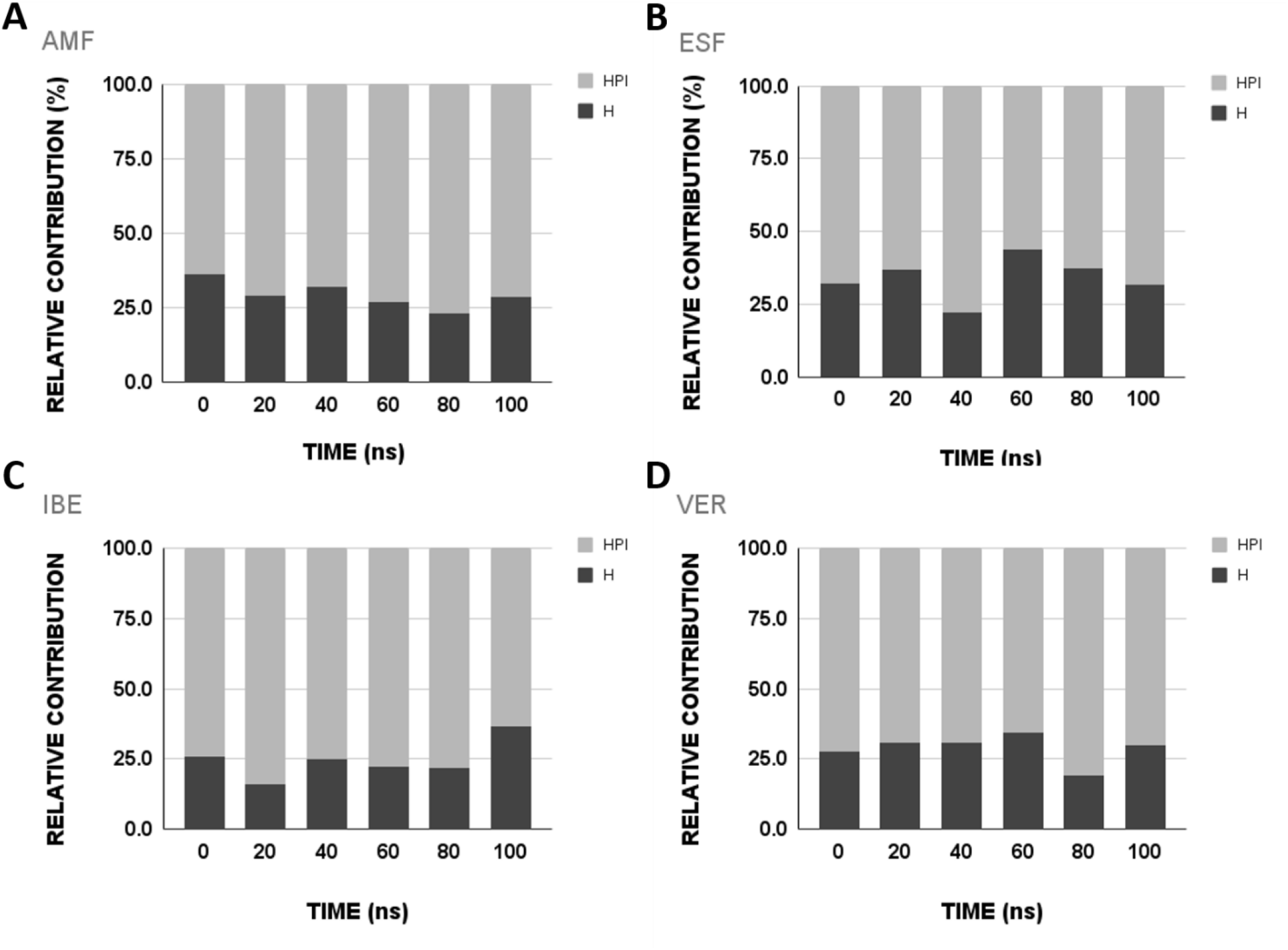
The relative contribution of hydrophobic interaction (HPI) and hydrogen bond (H BOND) in protein-inhibitor interaction of MD simulation presented in stacked bar diagram.

**Figure 6:**
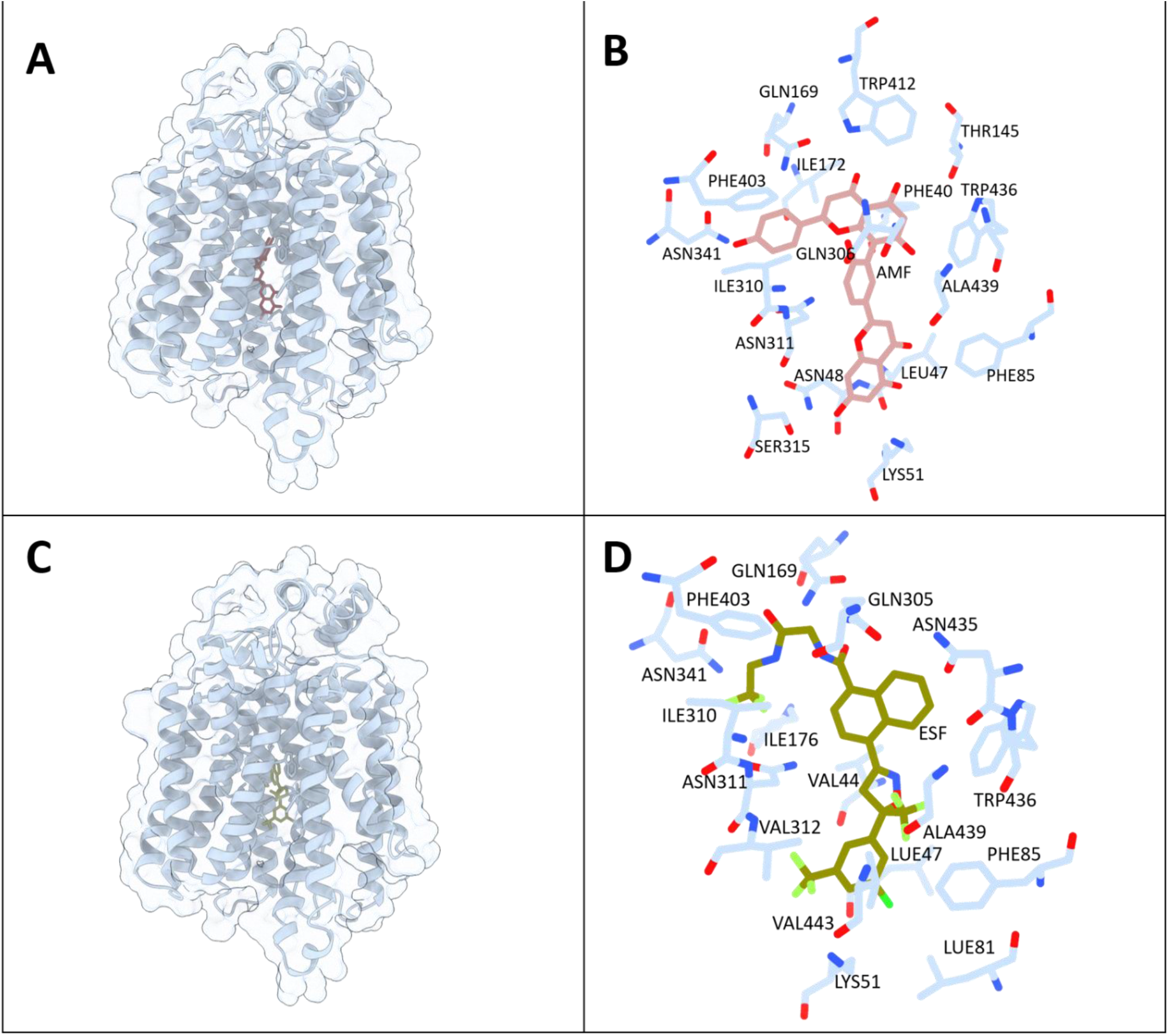
(A) The binding pose of AMF with respect to PfHT1 (B) residue level interaction between AMF and PfHT1 (C) The binding pose of ESF with respect to PfHT1 (D) residue level interaction between ESF and PfHT1. AMF, ESF and the interacting residues of protein are shown in sticks and labelled with the three-letter code.

### Rational Drug Design

A meticulous approach was employed for structure-guided de novo drug design, focusing on amino acid residue-level interactions between top inhibitor molecules and the target protein. From simulation metrics, as well as, binding energy calculated by MMPBSA, AMF and ESF were found to be highly stable, therefore these two molecules were utilized as scaffold to perform a structure guided drug design. Both AMF and ESF were fragmented into constituent functional groups and these were docked to PfHT1 to determine if they had specific affinity for any region of the binding pocket of the protein. Multiple cycles of design, modification and docking provided with three compounds, DH2, DH3 and DH4 with high affinity and specificity for PfHT1. A schematic presentation of drug designing strategy of DH2, DH3 and DH4 from AMF and ESF has been illustrated in Figure 7. The ADV energy of these three de-novo designed compounds along with their chemical structure are tabulated in Figure 8. The synthetic accessibility of these compounds along with ADME and toxicity score are presented in Supplementary Table 1. The Synthetic accessibility score for DH2, DH3, and DH4, is 4.11, 3.84 and 4.2 respectively, where 1 indicates very easy to synthesize and 10 indicates very difficult.

**Figure 7:**
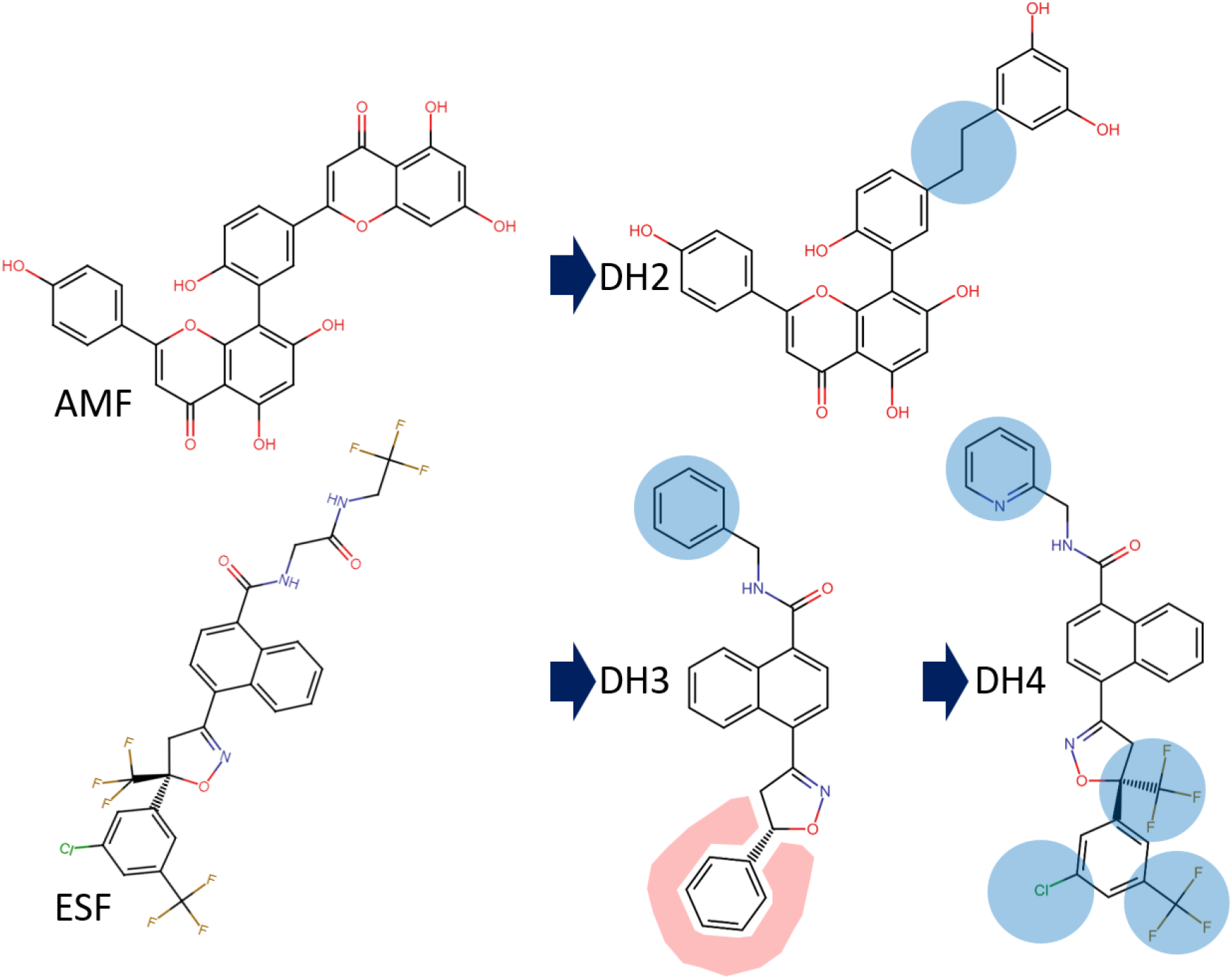
De novo drug design strategy. Where mother compounds, AMF and ESF are on left and the designed inhibitors, DH2, DH3, DH4 are on right, the site of modifications are marked with blue halos for addition and red halos for deletion.

**Figure 8:**
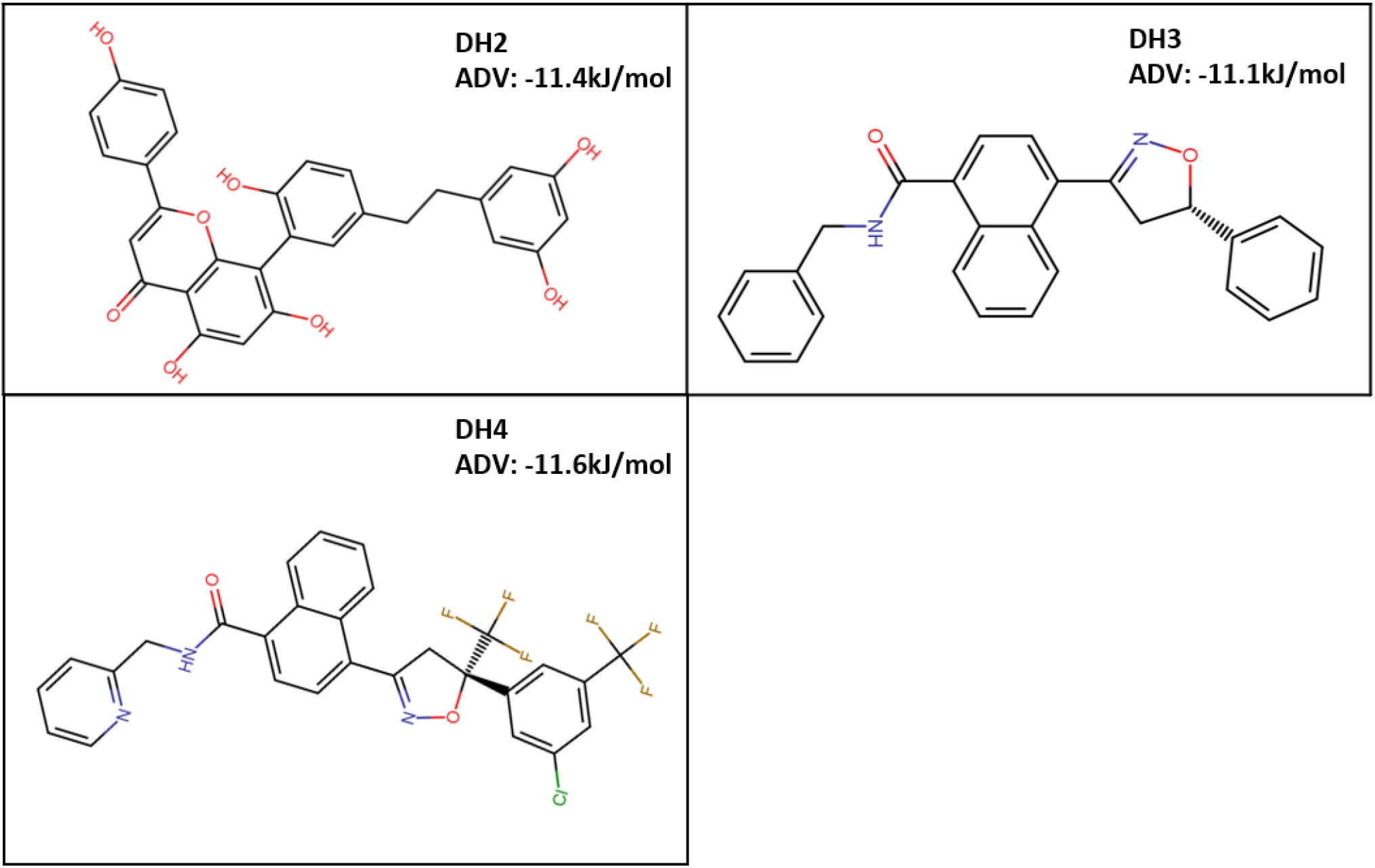
Name, chemical structure, the three-letter code and ADV energy of DH2, DH3, DH4.

The binding of these de novo designed compounds with the target protein was further validated by molecular dynamics simulation and MM-PBSA. Figure 9 represents the binding pose of DH2, DH3, and DH4 in the binding pocket of PfHT1 and also reveals the atomic level interaction between receptor and the designed molecules. The average RMSD, RMSF, SASA and Rg calculated from 100 ns MD simulation run are tabulated in Supplementary Table 2. The relative contribution of H bond and hydrophobic interaction in the binding of DH2, DH3 and DH4 are presented in Supplementary Figure 1. The dynamic nature of the protein-ligand complexes throughout the 100 ns time scale of MD run along with the conformational flexibility of amino acids are represented graphically in Figure 10. Binding free energies (ΔG) for DH2, DH3 and DH4 are calculated via MMPBSA method using last 20 ns trajectory of each MD run as described previously. Three of the de novo designed molecules showed highly favourable binding <-100 kJ/mol (Figure 11A). The contribution of Lennard-Jones potential and Coulombic short-range energy in the interaction of receptor-ligand is illustrated in Figure 11B. Figure 11C depicts ligand RMSD data, extracted from the 100 ns trajectories of MD simulation and compares the performance of the designed molecules, DH2, DH3 and DH4 with AMF. It clearly shows that in terms of ligand RMSD, DH4 shows best stability and the data is comparable with AMF, followed by DH2 whereas DH3 performed rather poorly. Considering ADV energy, ligand RMSD, binding free energy (ΔG) and other simulation data analysis parameters, out of 4500 compounds AMF has the potential to be considered a new lead molecule for structure-based drug designing as well as from the segment of de novo drug designing, DH4 shows promising results to be a highly specific inhibitor for PfHT1.

**Figure 9:**
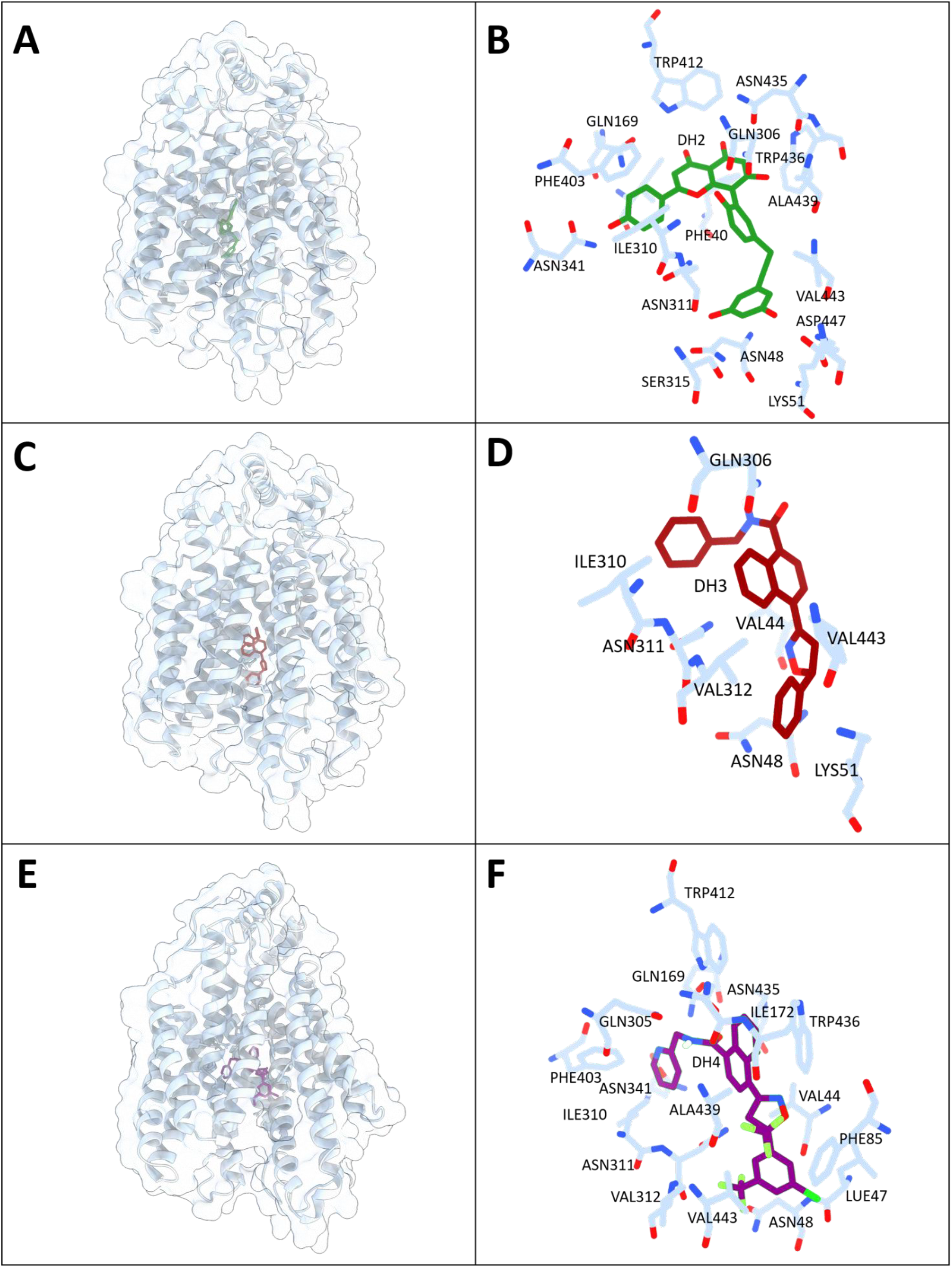
The binding pose and the interacting residues of DH2 (A,B), DH3 (C,D), DH4 (E,F). Inhibitors and the interacting residues of protein are shown in sticks and labelled with the three-letter code.

**Figure 10:**
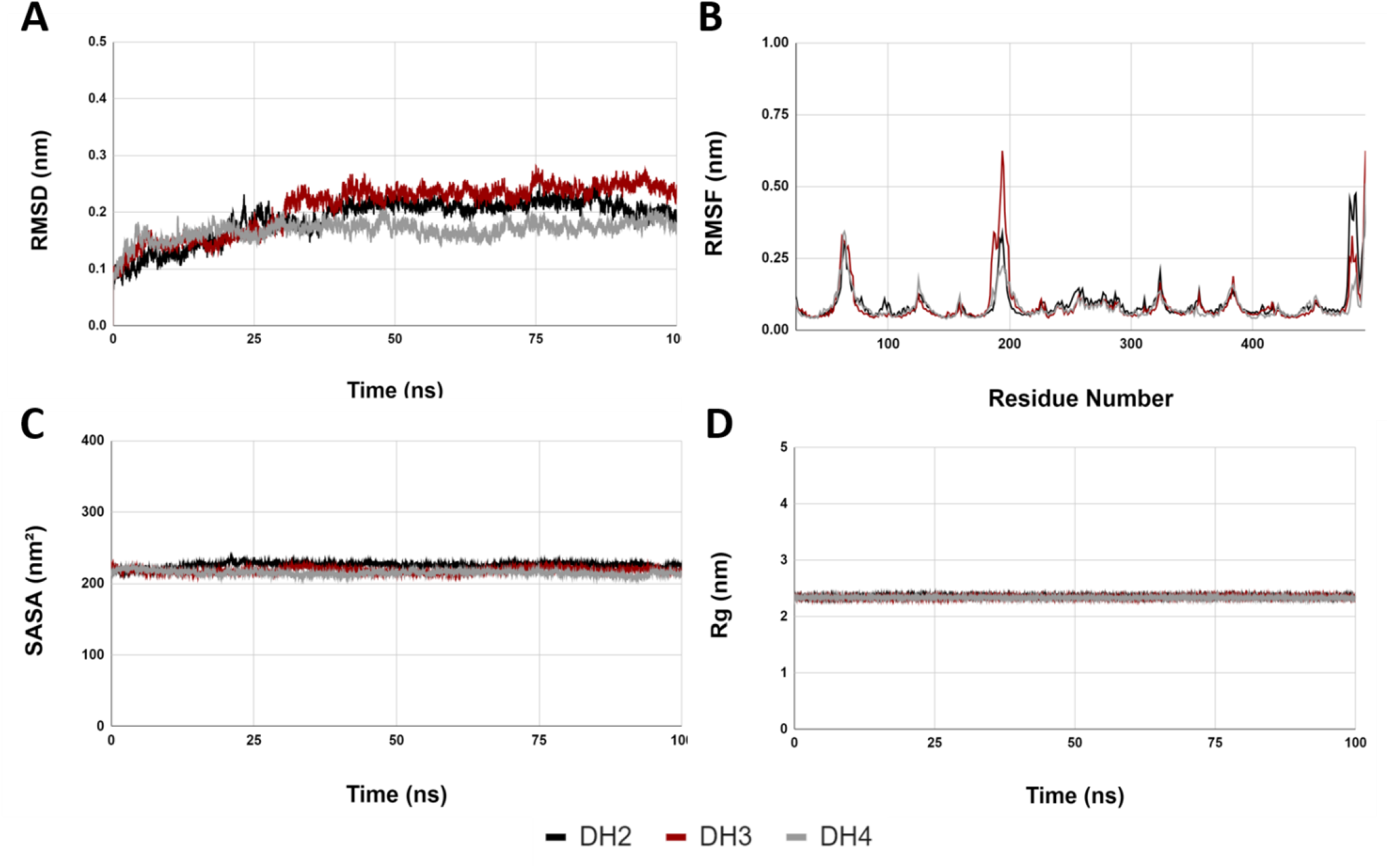
Molecular dynamics metrics of PfHT1 with de novo designed inhibitors, DH2, DH3, DH4. (A) The RMSD (nm) of backbone vs time in ns (B) backbone RMSF (nm) against the amino residue number, (C) solvent accessible surface area (SASA) in nm2 vs time (ns) and (D) Radius of gyration (Rg) in nm vs time in ns.

**Figure 11:**
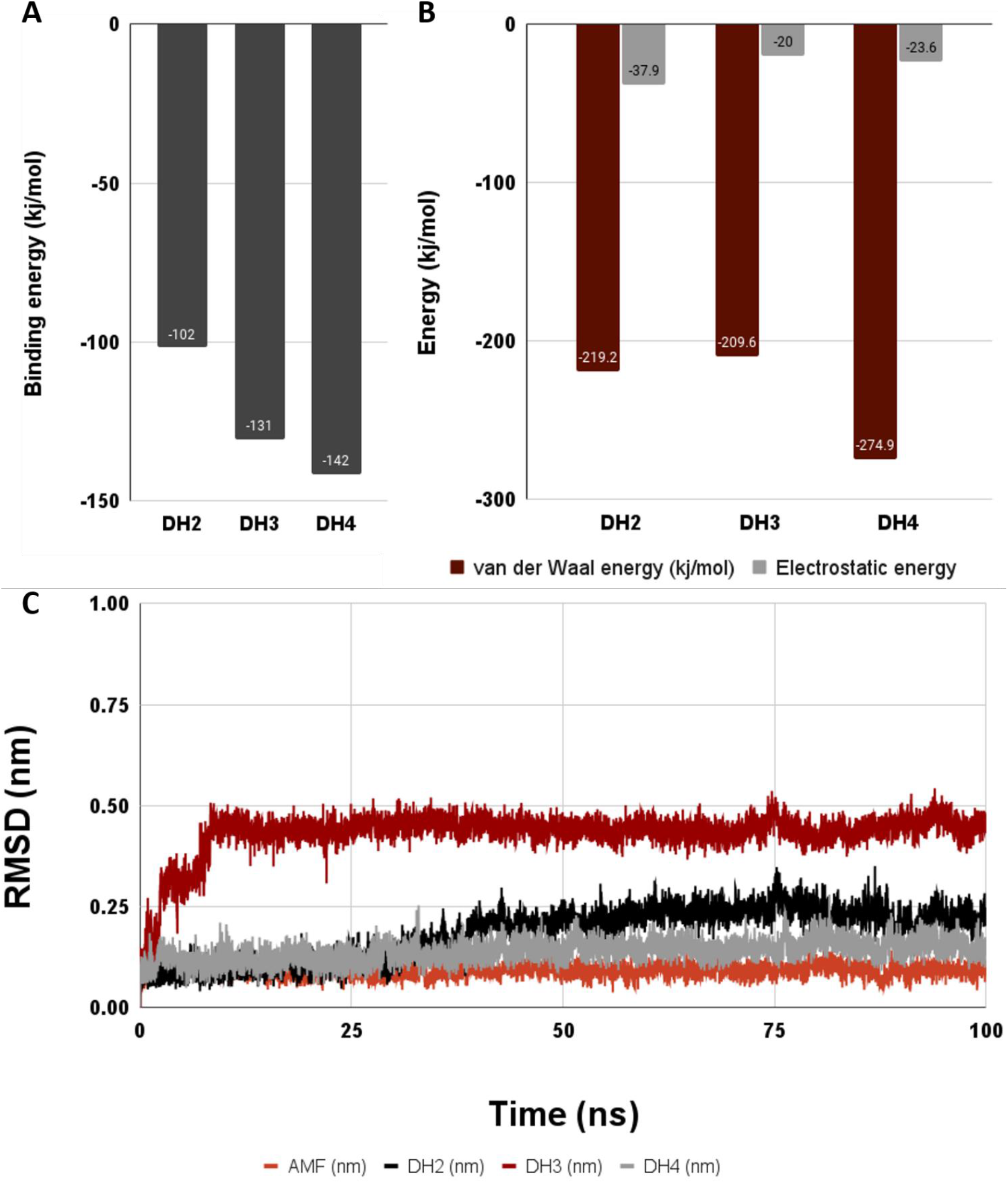
(A) Binding free energy (ΔG) for top 4, PfHT1-inhibitor complex expressed in kJ/mol. (B) The relative contribution of van der Waal’s energy and electrostatic energy in binding (expressed in kJ/mol) (C) Ligand RMSD calculated from simulation trajectory

Further analysis of PfHT1-AMF and PfHT1-DH4 interaction using PfHT1 crystal structures and MD simulation data from both apo and holo form of PfHT1 reveals that ligand binding triggers major conformational changes in the receptor. Comparing ligand bound structures with the apo form of PfHT1 by structural superposition shows that the side chain of PHE403 moved 2.52 Å towards AMF to form a stable stacking interaction where as TRP412 and PHE40 moved away 5.30 Å and 1.28 Å respectively from its native position in the apo form avoiding steric clash with the ligand (Figure 12A). Similarly, DH4 binding shifts the positions of PHE403 and ILE310, 4.67 Å and 7.31 Å respectively from its apo state to accommodate ligand in the binding pocket (Figure 12A). Besides such changes in amino acid side chain conformation at the ligand binding pocket, PfHT1 undergoes large conformation changes as depicted in Figure 12C, the loop spanning GLY183 to THR199 undergoes a conformational change of 13.06 Å in the ligand bound state. The helix loop helix structure spanning ASN52 to ILE75 undergoes approximately 50° bending motion towards central core of the transporter protein in the ligand bound state (Figure 12D). Which clearly indicates an open-close structural transition coupled with ligand binding. Figure 12E clearly shows that the helix loop helix structure is acting like a lid which keeps the transporter closed in the ligand bound state.

**Figure 12:**
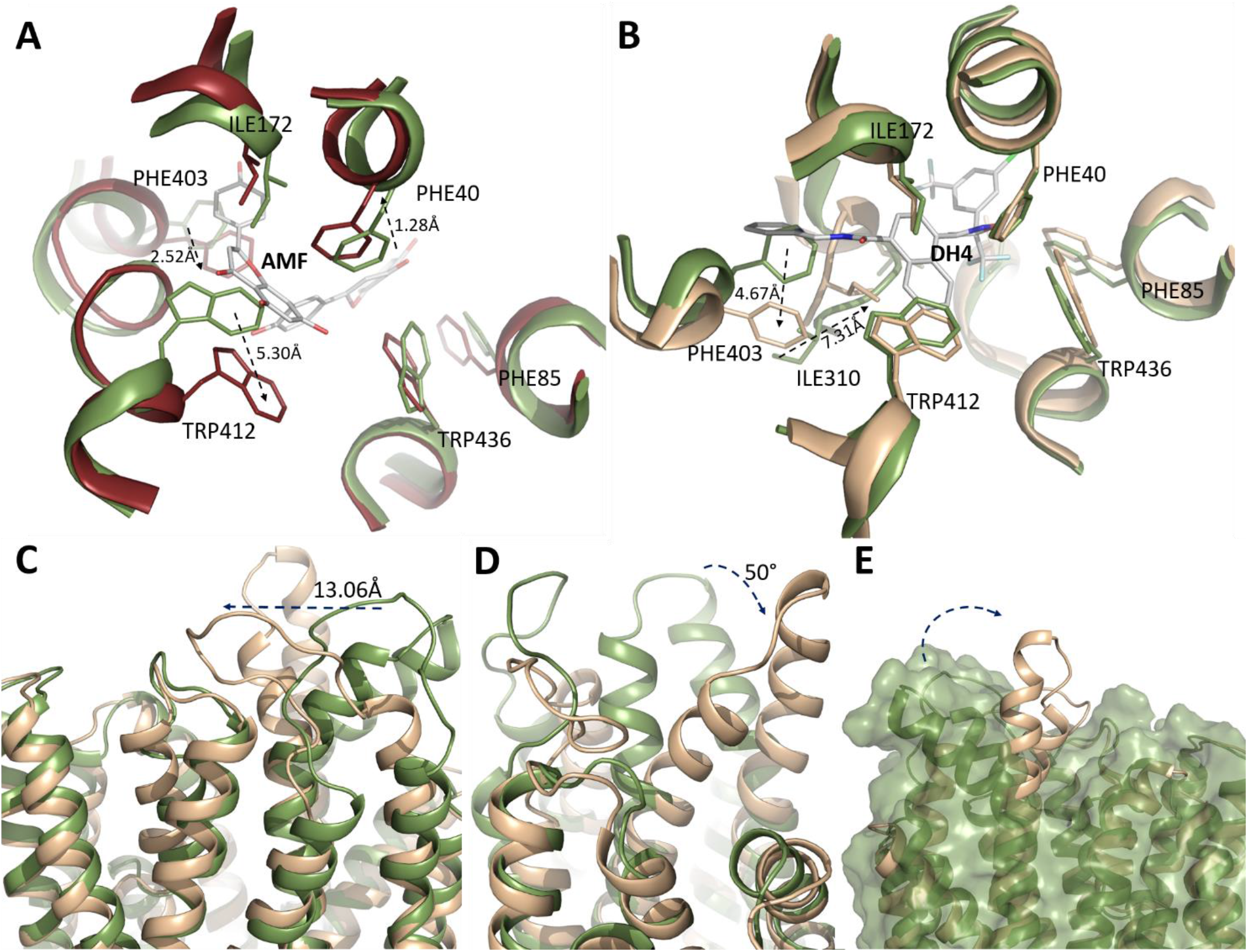
Conformational change. Comparison of spatial change of the amino acid side chains in the inhibitor binding pocket, with respect to AMF (A) and DH4 (B), where apo -protein is in green, AMF bound PfHT1 in ruby and DH4 bound PfHT1 in pale yellow. (C) Large conformational change that causes inhibitor induced closed state (pale yellow) when compared with the apo open form (green).

## CONCLUSION

The aim of this entire study was deciphering the mechanism of PfHT1-ligand interaction and the role of conformational dynamics of the PfHT1 in the process of transport. Using a wide variety of small molecules with different chemical nature coupled with molecular dynamics simulation revealed clear pattern of ligand induced large conformational change from open to close form. The conformational changes are also associated with reorientation of the amino acid side chains of the binding pocket at the central core of the protein. Authors carefully designed plasma membrane like environment with inner and outer polar atmosphere to perform all the simulations for not only to study the mechanism as well as finding novel inhibitors of PfHT1 via brute force screening method and selected two compounds, AMF & ESF out of 4500 compounds which passed all the validation tests. A unique strategy was adopted in this study where the chemical skeletons of AMF and ESF obtained from brute force screening were subjected to further modifications by de novo drug designing strategy. Of the de novo designed compounds, DH2, DH3 and DH4 was selected based on ADV energy, ease of synthesis and MD simulation metrics. DH4 showed highly stable ligand RMSD and best binding free energy, calculated using MMPBSA.

## MATERIALS AND METHODS

### Molecular Docking and Brute Force Screening

For brute force screening, Autodock Vina from PyRx platform (Dallakyan & Olson, 2015) facilitated high-throughput molecular docking. The X-ray crystal structure of *Plasmodium falciparum* hexose transporter, PFHT1 in complex with the C3361 inhibitor (PDB ID: 6M2L), retrieved from the Protein Data Bank (https://www.rcsb.org/). The missing residues were corrected by a template guided modelling on SwissModel (Bienert et al., 2017; Guex et al., 2009; Waterhouse et al., 2018). This modelled receptor was used for docking and simulation studies. The grid box parameters were set with a centre at coordinates -25.2, -45.9, 3.1, with dimensions 12.6 × 9.2 × 15.2. A library of over 4487 commercially available drug molecules in PDB format underwent preliminary screening, curated from diverse databases (e.g., PUBCHEM, Zinc, EMBL).

Screening and ranking were performed based on ADV docking scores, representing binding energy. The negative binding energy values, expressed in kcal/mol, indicated favourable binding between PfHT1 and small molecule ligands. The binding poses of selected small molecules exhibiting strong interactions with PfHT1 were validated using crystal structures of PfHT1 in both C3361-bound and glucose-bound forms.

Receptor and ligand preparation involved eliminating lone pairs, solvents, crystallization artifacts, and addressing missing hydrogen and charge, following a protocol detailed in previous work (Ganguly et al., 2023; Manna et al., 2023; Mitra et al., 2023).

### Drug-Like Properties and Toxicity Prediction

Following brute force screening, the top 4 compounds were subjected to a comprehensive analysis of Absorption, Distribution, Metabolism, and Excretion (ADME) along with toxicity prediction. This evaluation aimed to assess drug-likeness and safety profiles using the SwissADME server for ADME analysis and ProToxII for toxicity prediction.

In ADME analysis, adherence to Lipinski’s rule was determined based on five key physicochemical parameters: H-bond acceptors (NHBA), number of H-bond donors (NHBD), molecular weight, molar refractivity, and n-octanol/water partition coefficient. Compounds meeting these criteria, indicating favourable drug-like properties, were selected for further analysis.

The selected compounds, deemed druggable with low toxicity, underwent Molecular Dynamics Simulation (MD simulation). This step enhances our understanding of the compounds’ dynamic behaviour and stability over time, contributing valuable insights into their potential as drug candidates.

### Molecular Dynamics Simulation and MMPBSA Calculation Protocol

The PfHT1-inhibitor complexes were built with the CHARMM-GUI membrane builder (Jo et al., 2007, 2008; Park et al., 2021; Wu et al., 2014). To build the membrane bilayer, the receptor was inserted in a lipid bilayer comprised of 200 molecules of 1-palmitoyl-2-oleoylsn-glycero-3-phosphocholine (POPC) with a distribution ratio of 1:4 between the upper and the lower layer. The lipid bilayer was solvated in a water layer of 22.5Å in 150mM NaCl. The system comprising protein-ligand complex in a lipid bilayer was subjected to energy minimization, system equilibration (NVT and NPT), and MD run for 100ns. The simulation was performed using Gromacs 5.1.2(Lemkul, 2019), employing the Charmm36 all-atom force field (Brooks et al., 2009; Huang et al., 2017; Vanommeslaeghe et al., 2009).

RMSD (Schreiner et al., 2012), RMSF (Maiorov & Crippen, 1994), Rg (Lobanov et al., 2008), and SASA (Lemkul, 2019) computations provided comprehensive insights into the system’s dynamics. For binding free energy calculation, the final 20 ns trajectories were utilized. MMPBSA method was employed to compute the binding free energy (ΔG) in kJ/mol.

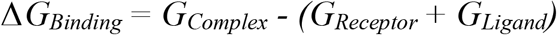

Selected drug molecules, undergoing MD simulation and MMPBSA-based binding free energy calculation, adhered to a stringent scientific protocol, enhancing the reliability and robustness of the study’s findings.

### Structure guided rational drug design

Molecules were designed using Chemical Sketch Tool from RCSB (RCSB PDB: Chemical Sketch Tool) the chemical skeleton underwent iterative modifications, validated by docking, targeting specific interaction functional groups.

## Acknowledgement

DH acknowledges St. Xavier’s College, Kolkata, India for providing necessary support. MG acknowledges WB-DST for the fellowship. AR acknowledges IIEST Shibpur for providing infrastructure and support.

## Contributions

All authors contributed to the study. All authors read and approved the final manuscript. DH and AR conceptualized the project, design the experiments, analyse the data, prepared the figures and wrote the manuscript. MG performed docking and MD simulation. MG and HM tabulated the results, performed preliminary analysis and helped in manuscript writing.

## Supplementary Data

**Supplementary Table 1:**
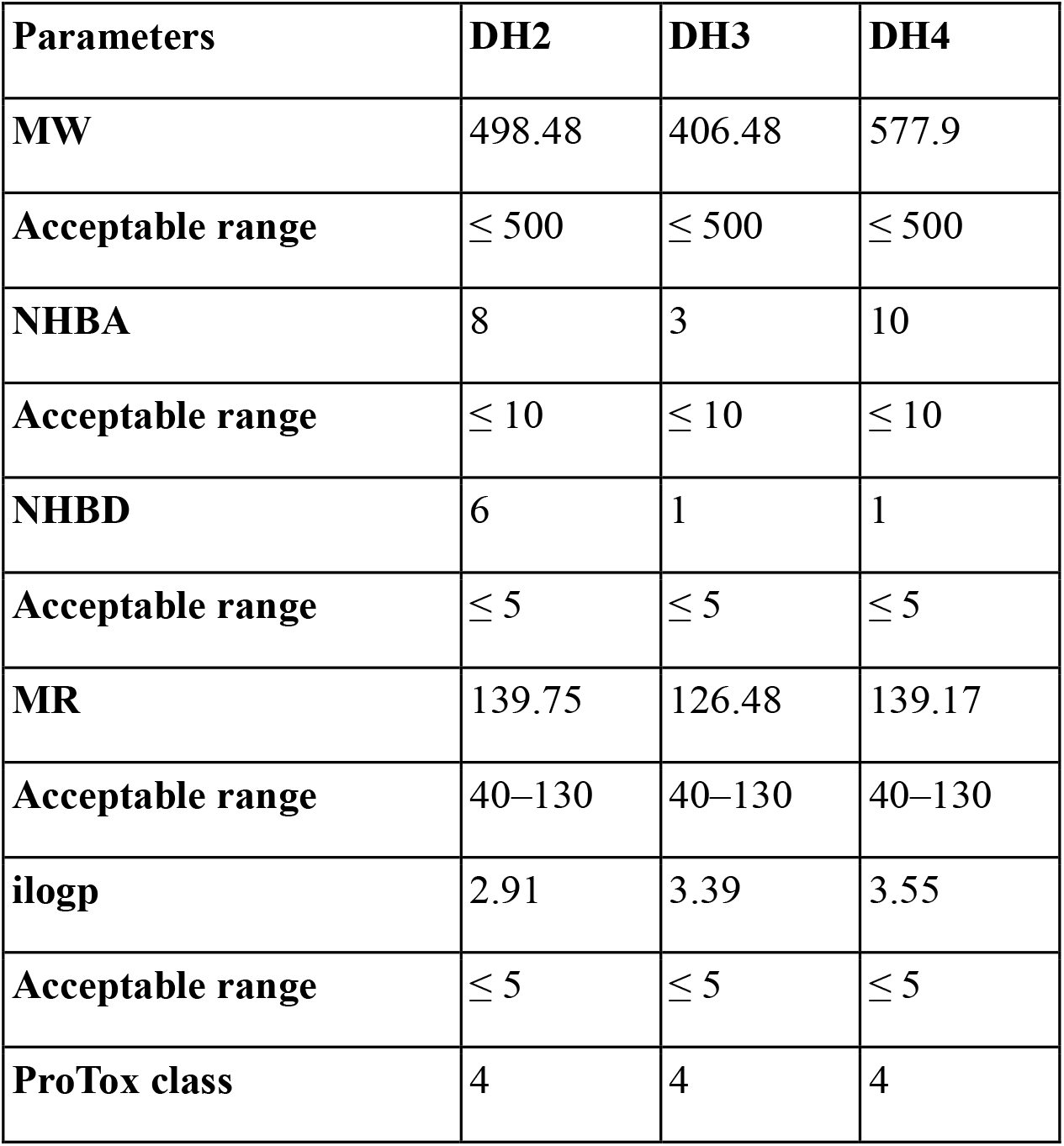
ADME and Toxicity assessment for DH2, DH3, DH4.

| Parameters | DH2 | DH3 | DH4 |
| --- | --- | --- | --- |
| MW | 498.48 | 406.48 | 577.9 |
| Acceptable range | $\leq 500$ | $\leq 500$ | $\leq 500$ |
| NHBA | 8 | 3 | 10 |
| Acceptable range | $\leq 10$ | $\leq 10$ | $\leq 10$ |
| NHBD | 6 | 1 | 1 |
| Acceptable range | $\leq 5$ | $\leq 5$ | $\leq 5$ |
| MR | 139.75 | 126.48 | 139.17 |
| Acceptable range | 40–130 | 40–130 | 40–130 |
| ilogp | 2.91 | 3.39 | 3.55 |
| Acceptable range | $\leq 5$ | $\leq 5$ | $\leq 5$ |
| ProTox class | 4 | 4 | 4 |

**Supplementary Table 2:**
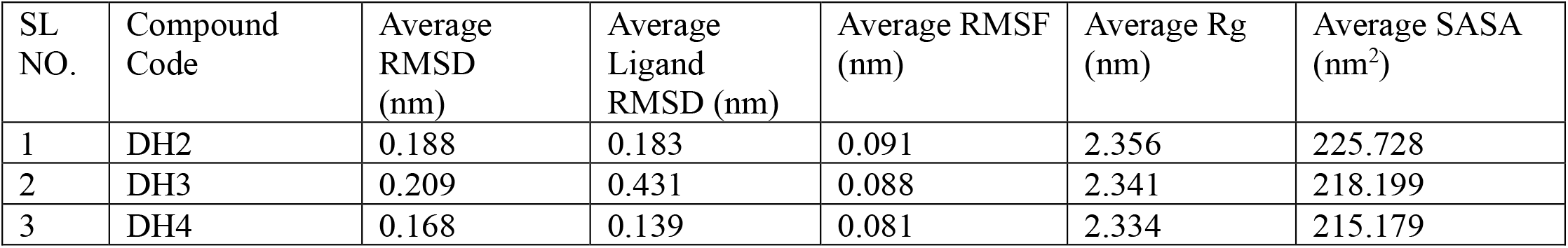
MD simulation metrics.

**Supplementary Figure 1:**
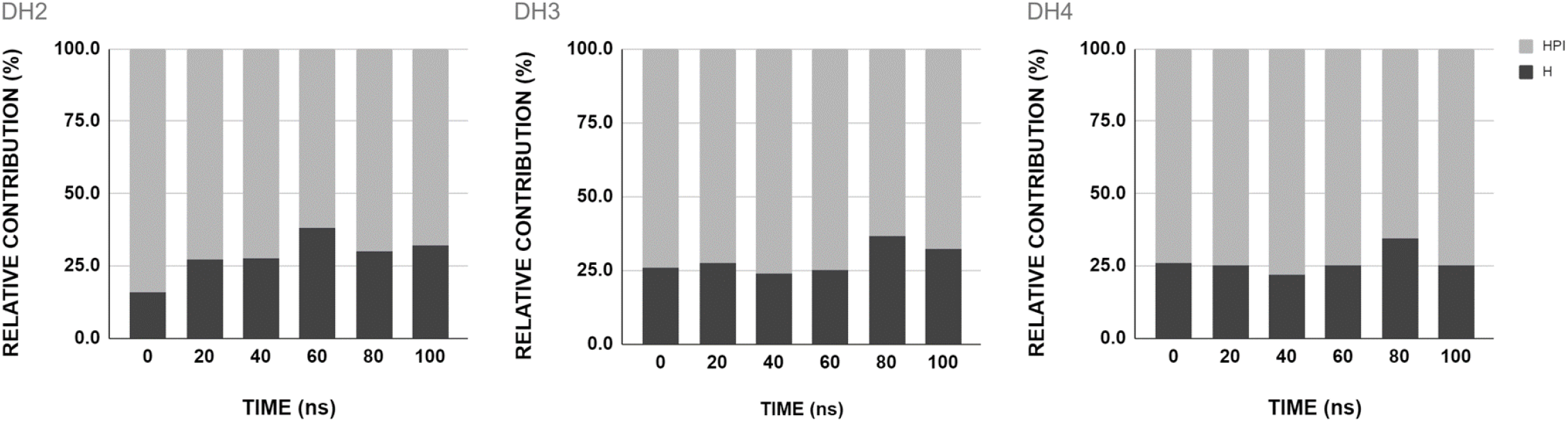
The relative contribution of hydrophobic interaction (HPI) and hydrogen bond (H BOND) in protein-inhibitor interaction of MD simulation presented in stacked bar diagram.

